# Longitudinal dynamics of clonal hematopoiesis identifies gene-specific fitness effects

**DOI:** 10.1101/2021.05.27.446006

**Authors:** Neil A. Robertson, Eric Latorre-Crespo, Maria Terradas-Terradas, Alison C. Purcell, Benjamin J Livesey, Joseph A. Marsh, Lee Murphy, Angie Fawkes, Louise MacGillivray, Mhairi Copland, Riccardo E. Marioni, Sarah E. Harris, Simon R. Cox, Ian J. Deary, Linus J. Schumacher, Kristina Kirschner, Tamir Chandra

**Affiliations:** MRC Human Genetics Unit, University of Edinburgh, Edinburgh, EH4 2XU, UK; Institute of Cancer Sciences, University of Glasgow, Glasgow, G61 1BD, UK; Cancer Research UK Beatson Institute, Glasgow, UK; Edinburgh Clinical Research Facility, University of Edinburgh, Edinburgh, EH4 2XU, UK; Centre for Genomic and Experimental Medicine, Institute of Genetics and Molecular Medicine, University of Edinburgh, Edinburgh, EH4 2XU, UK; Lothian Birth Cohorts, Department of Psychology, The University of Edinburgh, Edinburgh, UK; Department of Psychology, The University of Edinburgh, Edinburgh, UK; Centre for Regenerative Medicine, University of Edinburgh, Edinburgh, EH16 4UU, UK

## Abstract

The prevalence of clonal haematopoiesis of indeterminate potential (CHIP) in healthy individuals increases rapidly from age 60 onwards and has been associated with increased risk for malignancy, heart disease and ischemic stroke. CHIP is driven by somatic mutations in stem cells that are also drivers of myeloid malignancies. Since mutations in stem cells often drive leukaemia, we hypothesised that stem cell fitness substantially contributes to transformation from CHIP to leukaemia. Stem cell fitness is defined as the proliferative advantage over cells carrying no or only neutral mutations. It is currently unknown whether mutations in different CHIP genes lead to distinct fitness advantages that could form the basis for patient stratification. We set out to quantify the fitness effects of CHIP drivers over a 12 year timespan in older age, using longitudinal error-corrected sequencing data. We developed a new method based on drift-induced fluctuation (DIF) filtering to extract fitness effects from longitudinal data, and thus quantify the growth potential of variants within each individual. Our approach discriminates naturally drifting populations of cells and faster growing clones, while taking into account individual mutational context. We show that gene-specific fitness differences can outweigh inter-individual variation and therefore could form the basis for personalised clinical management.

## Introduction

Age is the single largest factor underlying the onset of many cancers (de Magalhaes, 2013). Age-related accumulation and clonal expansion of cancer-associated somatic mutations in healthy tissues has been posited recently as a pre-malignant status consistent with the multistage model of carcinogenesis (Martincorena, 2019). However, the widespread presence of cancer-associated mutations in healthy tissues highlights the complexity of early detection and diagnosis of cancer (Ayachi et al., 2020; Genovese et al., 2014; Jaiswal et al., 2014; Lee-Six et al., 2019; Martincorena et al., 2015).

Clonal haematopoiesis of indeterminate potential (CHIP) is defined as the clonal expansion of haematopoietic stem and progenitor cells (HSPCs) in healthy aged individuals. CHIP affects more than 10% of individuals over the age of 60 years and is associated with an estimated 10-fold increased risk for the later onset of haematological neoplasms (Ayachi et al., 2020; Genovese et al., 2014; Jaiswal et al., 2014). There is a clear benefit of detecting CHIP early as the association between clone size and malignancy progression is well-established (Jaiswal and Ebert, 2019; Jaiswal et al., 2014; Park and Bejar, 2020).

The particular mechanisms by which common mutations of CHIP, e.g. *DNMT3A*, *TET2*, contribute to the progression of leukaemia are still not understood, which hinders early diagnosis of CHIP on a gene or variant-basis (Challen and Goodell, 2020; Jaiswal and Ebert, 2019; Shih et al., 2012; Terradas-Terradas et al., 2020). In clinical practice, CHIP is diagnosed by the presence of somatic mutations at variant allele frequencies (VAF) of at least 0.02 in cancer-associated genes in more than 4% of all blood cells (Jaiswal and Ebert, 2019; Steensma and Bolton, 2020). Clonal fitness, defined as the proliferative advantage of stem cells carrying a mutation over cells carrying no or only neutral mutations, has emerged as an alternative clone-specific quantitative marker of CHIP (Watson et al., 2020; Williams et al., 2020). As mutations in stem cells often drive leukaemia (Jaiswal et al., 2014), we hypothesise that stem cell fitness contributes substantially to transformation from CHIP to leukaemia.

Stratification of individuals to inform clinical management for early detection or prevention of leukaemia will depend on our ability to accurately associate genes and their variants with progression to disease. However, it remains unresolved whether variant- or gene-specific fitness effects transcend other factors contributing to variable progression between individuals such as environment or genetics.

Hitherto fitness effects have been predicted from large cross-sectional cohort data (Abelson et al., 2018; Watson et al., 2020). In this approach, single time-point data from many individuals is pooled to generate allele frequency distributions. Although this method allows the study of a large collection of variants, pooling prevents estimation of an individual’s mutational fitness effects from cross-sectional data. Inferring fitness from a single time-point creates additional uncertainty about whether a mutation has arisen recently and has grown rapidly (high fitness advantage), or arose a long time ago and grown slowly (low fitness advantage). With longitudinal samples, fitness effects of individual mutations can be estimated directly from the change in VAF over multiple time-points.

In this study we work with longitudinal data from the Lothian Birth Cohorts of 1921 (LBC1921) and 1936 (LBC1936). Such longitudinal data are rare worldwide owing to their participants’ older age (70-90 years) and their three-yearly follow-ups over 12 years in each cohort and over 21 years total timespan. We developed a new framework for extracting fitness effects from longitudinal data. Firstly, a drift-induced fluctuations (DIF) filter allowed us to better segregate between naturally drifting populations of cells and fast-growing clones. Secondly, a context-dependent fitting method quantified the growth potential or fitness effects simultaneously for all selected mutations within each individual. We detected gene-specific fitness effects within our cohorts, highlighting the potential for personalised clinical management.

## Material and Methods

### Participant samples

The Lothian Birth Cohort 1921 (LBC1921) contains a total of 550 participants at Wave 1 of their testing (done between 1999 and 2001) with a gender ratio of 234/316 (m/f) and a mean age at Wave 1 of 79.1 (SD=0.6) (Table 1, (Taylor et al., 2018)). The Lothian Birth Cohort 1936 (LBC1936), contains a total of 1091 participants at Wave1 of their testing (done between 2004 and 2007) with a gender ratio of 548/543 (m/f) and a mean age at Wave 1 of 69.5 (SD=0.8) (Table 1, (Taylor et al., 2018)). We previously identified 73 participants with CHIP at Wave 1 (Robertson et al., 2019). We sequenced DNA from those 73 LBC participants longitudinally and added 16 LBC participants with previously unidentified CHIP, for a total of 298 samples together with 14 “Genome in a Bottle” (GIAB) controls (Supplemental Table 1).

### Targeted, error-corrected Sequencing and Data filtering

DNA was extracted from EDTA whole blood using the Nucleon BACC3 kit, following the manufacturer’s instructions. Libraries were prepared from 200ng of each DNA sample using the Archer VariantPlex 75 Myeloid gene panel and VariantPlex Somatic Protocol for Illumina Sequencing (Invitae, Table 2), including modifications for detecting low allele frequencies. Sequencing of each pool was performed using the NextSeq 500/550 High-Output v2.5 (300 cycle) kit on the NextSeq 550 platform. To inform reproducibility, background model for error and batch correction, we sequenced “genome in a bottle” DNA in each batch of samples (DNA NA12878 Coriell Institute).

Reads were filtered for phred ≥30 and adapters removed using Trimmomatic (Bolger et al., 2014) before undergoing guided alignment to human genome assembly hg19 using bwa-mem and bowtie2. Unique molecular barcodes (ligated prior to PCR amplification) were utilised for read deduplication to support quantitative multiplexed analysis and confident mutation detection. Within targeted regions, variants were called using three tools (Lofreq (Wilm et al., 2012), Freebayes (Garrison and Marth, 2012) and Vision (ArcherDX, unpublished) – building a consensus from the output of all callers (Table 3).

All filtered variants at 0.02 VAF met the following criteria: 1) the number of reads supporting the alternative allele surpasses the coverage criteria while exhibiting no directional biases (AO ≥ 5, UAO ≥ 3); 2) variants are significantly underrepresented within the Genome Aggregation Database (gnomAD; p ≤ 0.05); 3) variants are not obviously germline variants (stable VAF across all waves ~0.5 or ~1) that may have been underrepresented in the gnomAD database due to the narrow geographical origin of the Lothian Birth Cohort participants; 4) contain events that are overrepresented across the dataset - generally frameshift duplications and deletions - whose reads share some sequence homology to target regions yet are likely misaligned artefact from the capture method. In addition, we manually curated this list checking for variants that were previously reported, as per Jaiswal et al. (Jaiswal et al., 2014b), in the COSMIC (Catalogue of Somatic Mutations in Cancer) database or within the published literature (Table 4). Finally, for any variant that surpassed the above criteria at, or above, 0.02 VAF across the measured time period, we endeavored to include all other participant matched data points regardless of VAF level (SuppFig.1A).

### Computational prediction of missense variant effects

To predict which missense variants are most likely to be damaging, we used six computational variant effect predictors recently identified as being most useful for identifying pathogenic mutations (Carter et al., 2013; Hecht et al., 2015; Ioannidis et al., 2016; Livesey and Marsh, 2020; Raimondi et al., 2017; Riesselman et al., 2018; Vaser et al., 2016). Specifically, for each variant identified in this study, we determined what fraction of previously identified pathogenic and likely pathogenic missense variants from ClinVar, and what fraction of variants observed in the human population from gnomAD v2.1 for each computational predictor. We then averaged these fractions across all predictors. Note that DeepSequence was only included for DNMT3A and TP53 due to its computational intensiveness and difficulty of running on long protein sequences.

### Modelling to estimate fitness

Given the longitudinal nature of this study we can directly fit the deterministic solution of an established minimal model of cell division (Till et al., 1964; Watson et al., 2020) to the time evolution of VAF trajectories in a participant’s genetic profiles (Fig.2C). For each individual we simultaneously estimate the fitness and time of acquisition of variants as well as the size of the stem cell pool.

In this model cells exist in two states: stem cells (SCs) or differentiated cells (DCs). Under the assumption that DCs cannot revert to a SC state, differentiation inevitably leads to cell death and is treated as such. Furthermore, assuming that each SC produces the same amount of fully differentiated blood cells allows a direct comparison between the VAF of a variant as observed in blood samples and the number of SCs forming the genetic clone (clone size). For an individual with a collection of clones {*c_i_*}_*i∈I*_, the VAF evolution in time *v_i_*(*t*) of a clone *c_i_* corresponds to 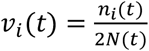, where *v_i_*(*t*) is the variant allele frequency of the variant at time *t*, *n_i_*(*t*) is the number of SCs carrying the variant and *N(t)* corresponds to the total number of diploid HSPCs present in the individual. Finally, we assume that that *N(t)* = *N_w_* + ∑_*i∈I*_ *n_i_*(*t*) where *N_w_* is the average number of wildtype (WT) HSPCs in the individual.

The bias towards self-renewal of symmetric divisions is parameterised by *s* and determines the fitness advantage of a clone. In normal haematopoiesis *s* = 0, in which case clones undergo neutral drift. The stochastic time-evolution of neutral clones corresponds to that of a critical birth and death model (CBM) that we exploit to identify mutations with a clear proliferative advantage, as detailed below. For clones with non-neutral (fitness-increasing) mutations, *s* > 0, and these clones grow in size exponentially in time as 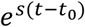 from an initial population of 1 SC at the time of mutation acquisition *t*_0_.

Altogether, the exponential proliferation of an isolated genetic clone in an individual results in the logistic time-evolution of its corresponding VAF,

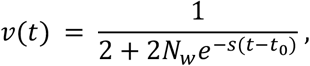

allowing us to estimate the fitness advantage in trajectories with at least 3 data points - 46% of all fit trajectories - using least-squares fitting. Further, when multiple fit clones are present in an individual we constrain the fit to share the stem cell pool size *N(t)* for all variant trajectories in this individual. This increases the data/parameter ratio, and produces richer dynamics, where exponentially growing clones can be suppressed by the growth of a fitter clone. Critically, this implies that even non-competitive models, where trajectories grow independently of each other, will result in competitive dynamics in the observed VAF trajectories as variants strive for dominance of the total production of blood cells.

Notice that this model cannot account for loss of heterozygosity events. Overall, only one variant - *JAK2 c.1849G>T* - averaged a VAF > 0.5 and was consequently left out of this study. Further, following the fit, three filters were implemented to detect inaccurately fitted trajectories left out from the reported fitness estimates: 1) individuals presenting a large r^2^-value, total squared error accumulated over all fitted trajectories in a participant (2 of 54 participants); 2) trajectories with a negative Pearson correlation coefficient between fit and observed data, e.g. a growing trajectory with a decreasing fit, which could result from fitter variants that were not captured in our gene panel (34 of 90 trajectories when using a 0.02 VAF filter, 8 of 96 trajectories when using a drift-induced fluctuations filter, see below and Results for filtering approaches); 3) individuals with clinical records indicating treatment that could affect the clonal dynamics (1 participant, see SI methods). Both exclusion criteria 1 and 2 included participants that likely had clones with multiple mutations based on visual inspection.

### Filtering fit variants using mathematical modelling of neutral drift

A critical birth-death process (Fig.2C) predicts drift-induced fluctuations (DIF) in the size of clones with zero mean and variance *σ*(*t*)^2^ = 2*λn*(*t*_0_)*t*, where *n*(*t*_0_) is the clone size at the start of the observation period of length *t*, and *λ* is the rate of symmetric self-renewing cell divisions. This results in a predicted linear relationship between the VAF of a clone, *v*, and the expected variance in the distribution of its fluctuations, *Δv* (VAF gradient between two time points normalised to be independent of time, SI methods). More precisely, the distribution of DIF, *Δv*, will have expectation *μ_Δv_* = 0 and variance *σ_Δv_*^2^ = *v*(*t*_0_)*λ*/*N*, where *N* is again the total number of HSPCs in the individual. In the presence of a sufficiently large growing non-neutral mutation, the expected size of DIF of neutral clones is no longer zero but decreases with VAF (SI methods), *μ_Δv_* ~ −*v*(*t*_0_), while *σ_Δv_*^2^ remains proportional to VAF (SI methods).

We use the longitudinal trajectories of synonymous mutations, as a proxy for neutral mutations, in the LBC to fit linear regressions *μ_Δv_* ~ −*v*(*t*_0_) and *σ_Δv_*^2^ ~ *v*(*t*_0_) (Fig.3B). The DIF filter then identifies variants whose growth cannot be explained by neutral drift fluctuations, that is variants *v* satisfying *Δv* > *μ_Δv_* + 2*σ*_*Δv*_(SI methods).

## Results

### The Lothian Birth Cohorts allow for longitudinal profiling of CHIP variants in advanced age

The Lothian Birth Cohorts (LBCs) of 1921 (n=550) and 1936 (n=1091) are two independent, longitudinal studies of ageing with approximately three yearly follow up for five waves, from the age of 70 (LBC1936) and 79 (LBC1921) (Taylor et al., 2018). We previously identified 73 participants with CHIP at Wave 1 through whole-genome sequencing (Robertson et al., 2019). Here, we used a targeted error-corrected sequencing approach using a 75 gene panel (ArcherDX/Invitae) to assess longitudinal changes in variant allele frequencies (VAF) and clonal evolution over 21 years across both LBC cohorts and 12 years within each cohort (Table 1). Error-corrected sequencing allowed accurate quantification, providing more sensitive clonal outgrowth estimates compared to our previous WGS data. We sequenced 298 samples (89 individuals across 2-5 time-points) and achieved a sequencing depth of 1953x mean coverage (1930x median) over all targeted sites with an average of 2.4 unique somatic variants (pan-cohort VAF 0.0005-0.87, median VAF 0.025) detected per participant. We examined all participant-matched events across the time-course: sequence quality control metrics revealed that only 12 of 466 data-points failed to meet our quality criteria likely due to low initial VAF. The majority of our variant loci generally displayed a high number of supporting reads, with a mean of 186 (SuppFig.1A).

For our initial analysis, we retained variants with at least one time-point at 0.02 VAF (Table 5). *DNMT3A* was the most commonly mutated CHIP gene (n=53 events in 42 participants), followed by *TET2* (n=22 events in 18 participants), *NOTCH1* (n=10 events in 9 participants), *JAK2* (n=8 events in 8 participants) and *U2AF2* (n= 7 events in 6 participants) (Fig.1A and 1C). Our mutation spectrum is consistent with previous studies in finding *DNMT3A* and *TET2* as the most frequently mutated genes. However, we detected a lower frequency of mutations in splicing genes, such as *SRSF2, U2AF1* and *SF3B1*, despite the older age of the cohort (Fig.1A and SuppFig.1A). This is in contrast to previously published cohort data, where splicing mutations became more prominent with increased age (McKerrell et al., 2015). The majority of mutations were missense, frameshift and nonsense mutations (Fig.1B). Clonal evolution and changes in VAF per participant over all time points are shown in Fig.1C and SuppFig.1C. We detected some variants more frequently at certain hot spots within a gene such as p.Arg882His in *DNMT3A*, with previously unreported variants being present as well (Fig.1D-I, Table 2, (Jaiswal et al., 2014)). In the case of *JAK2V617F*, we identified two individuals who developed leukaemia at Wave 2 and received treatment between Waves 2 and 3, likely driving a clear reduction in clone size (Fig.1H). Those indivdiuals were excluded from further analysis. Overall, our sequencing approach allowed for high resolution, longitudinal mapping of CHIP variants over a 12-year time span per cohort and 21 year timespan across both cohorts from the same geographical region and born 9 years apart.

**Figure 1.**
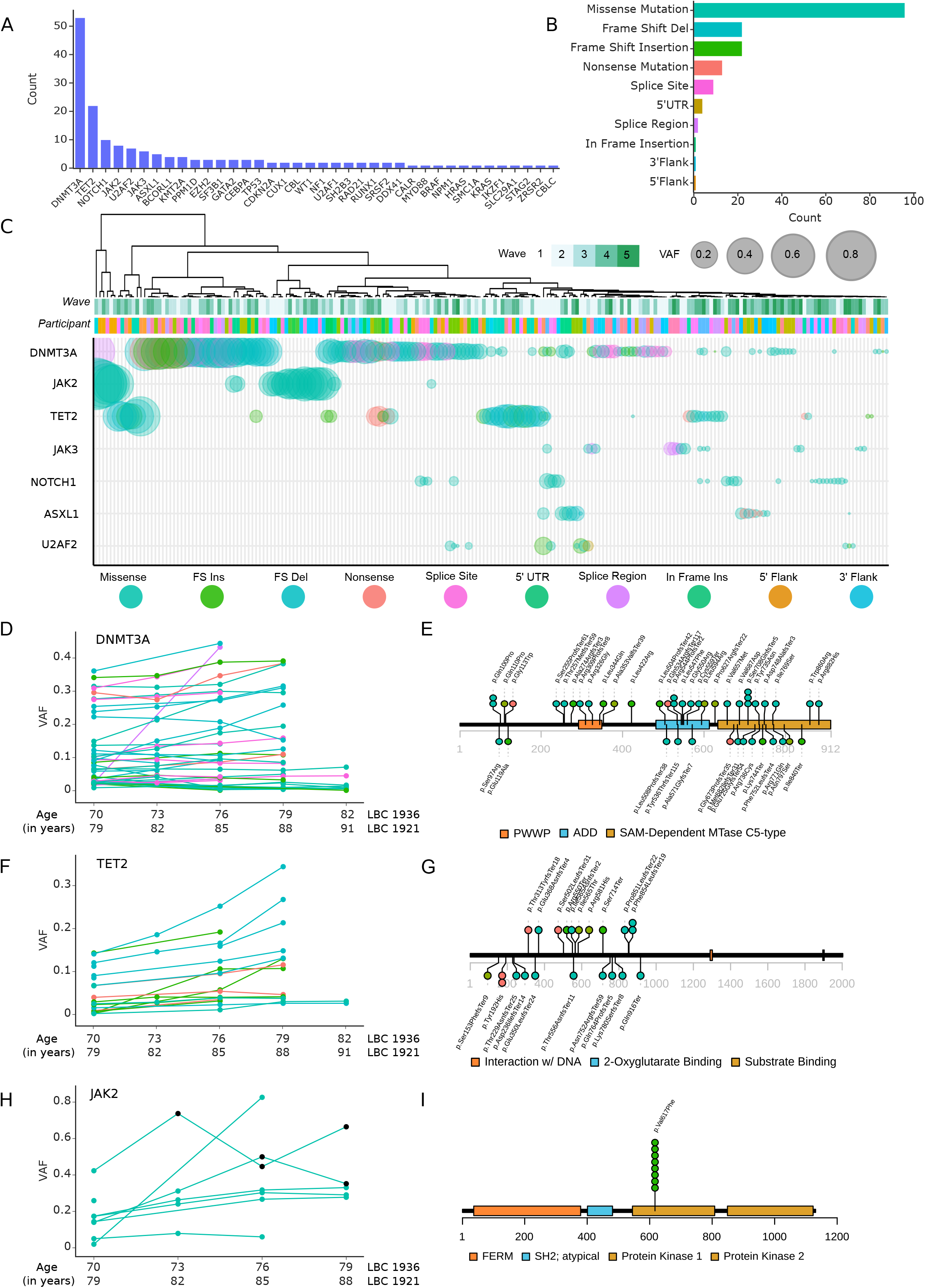
Clonal Haematopoiesis in the Lothian Birth Cohorts of 1921 and 1936. A. Counts of unique events that exceeded 0.02 VAF across the range of the longitudinal cohorts (approx. 70 - 79 years and 79 - 91 years in LBC36 and LBC21, respectively) in our panel of 75 haematopoietic genes. B. Counts of the functional consequences of the unique events listed in Fig.1A, highlighting missense mutations as the most frequently encountered event type. Several other key protein altering event types rank highly, including frameshift insertions/deletions and nonsense mutations. C. Schematic of the top seven most affected genes in the cohort with the largest clone size of an event in any given gene shown. All affected participants have been clustered across all time-points, with the point size scaled by VAF and coloured by the functional consequence of the variant (as per Fig.1B and legend). Participants broadly cluster together across their time-course, driven by the expanding or stable VAF of their harboured mutations and underscores the high prevalence and large clone size of common clonal haematopoietic drivers, namely, DNMT3A, TET2 and JAK2. D. Clone size trajectories of all DNMT3A mutations across the time-series in both LBC21 and LBC36 coloured by the functional consequence of the variant (as per Fig.1B and 1C). E. Locations of the somatic mutations discovered in DNMT3A. Protein affecting events are marked and labeled across the structure of the gene (missense in red, truncating in purple, stacked for multiple events) highlighting a strong, previously described tendency for CH events in DNMT3A to occur in conserved structural and domain loci (which are coloured and labeled along the amino acid length of the protein). F. Clone size trajectories of all TET2 mutations across the time-series in both LBC21 and LBC36 coloured by the functional consequence of the variant (as per Fig.1B and 1C). G. The locations of somatic mutations in TET2. Protein affecting events are marked and labeled across the structure of the protein (missense in red, truncating in purple, stacked for multiple events). H. Clone size trajectories of all JAK2 mutations across the time-series in both LBC21 and LBC36 coloured by the functional consequence of the variant (as per Fig.1B and 1C). Points marked in black denote time-points after which the affected participant received treatment for leukaemia - potentially driving the observed reductions in clone size. I. The locations of somatic mutations in JAK2. Protein affecting events are marked and labeled across the structure of the protein (missense in red, truncating in purple, stacked for multiple events). Here, all eight JAK2 mutations are p.Val617Phe (JAK2-V617F) missense variants.

### Identification of harmful CHIP variants in the LBCs

Stem cell fitness is defined as the proliferative advantage over cells carrying no or only neutral mutations. It remains incompletely understood to what extent fitness is gene- or variant-specific, or determined by the bone marrow microenvironment and clonal composition. Earlier estimates suggested a wide spread of fitness effects even for variants of the same gene (Watson et al., 2020), which would make it difficult to clinically stratify individuals with CHIP. To determine the fitness effects of the variants identified in our cohorts (Fig.1A), we initially selected all CHIP variants in our data using the commonly used criterion of defining any variants with VAF>0.02 as CHIP (Jaiswal and Ebert, 2019; Steensma and Bolton, 2020), and retaining only those variants with at least 2 time-points (Fig.2A). This approach identified 140 CHIP mutations (Fig.2B) overall. To estimate the fitness effect each variant confers, we fitted longitudinal trajectories using birth-death models of clonal dynamics (Fig.2C) on trajectories with 3 or more time-points (Table 6). The resulting fitness values show an overall dependence of fitness on the gene level (Fig.2E), with a wide distribution of fitness for some genes, such as TET2 and DNMT3A (Fig.2D and 2E), but not others such as JAK2 (which are all the same variant).

**Figure 2.**
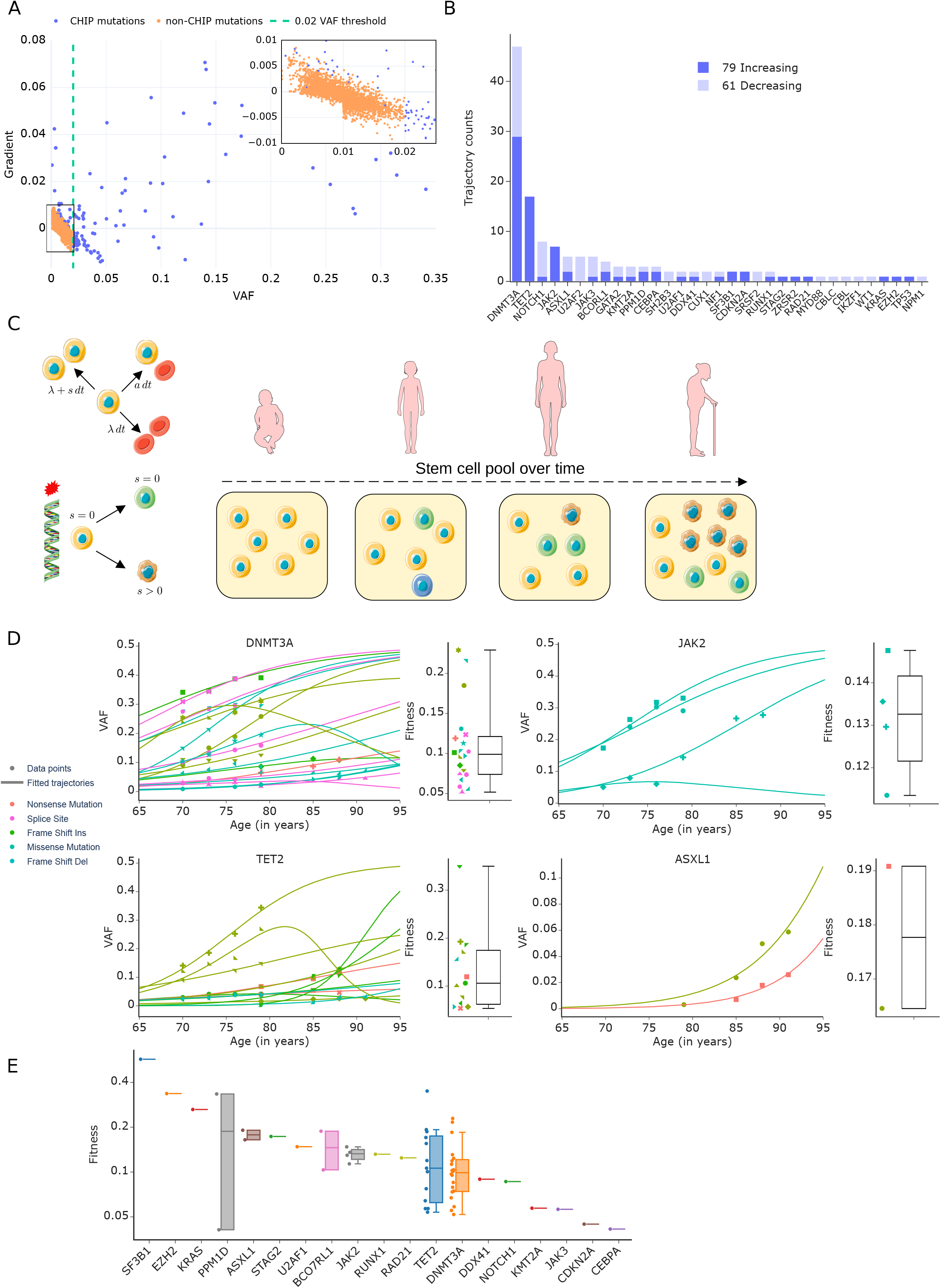
Fitness effects of variants thresholded at 0.02 VAF. A. VAF measurement *v*(*t*_0_) at initial time-point *t*_0_ vs time-normalized gradient in VAF, 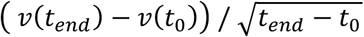, between initial and last time-points *t*_0_ and *t_end_* of all variants detected in the LBC with at least 2 time-points. Each data point corresponds to a trajectory in the LBC and has been coloured according to its CHIP status based on the 0.02 VAF threshold (dashed green line). Blue and orange respectively denote whether or not trajectories achieved a VAF > 0.02 during the observed time-span. Inset: Close-up of low VAF region showing a large number of variants below 0.02 VAF. Note data below 0.01 VAF were only included if the trajectory later rises above 0.01, hence the data are less dense below VAF=0.01. B. Number of trajectories passing the currently used 0.02 VAF-threshold, broken down into whether VAF is increasing or decreasing from first to last time-point. C. Schematic of mathematical model of clonal dynamics. Stem cells naturally acquire mutations over time leading to the formation of genetic clones. Artwork includes images by Servier Medical Art licensed under CC BY 3.0. D. Fitted trajectories (lines) and fitness estimates (box-plots) for all variants in selected genes DNMT3A, JAK2, ASXL1, TET2. Note decreasing trajectories occur even for non-zero fitness because our model fitting takes into account the presence of other fit mutations in the same individual. E. Fitness effects of mutations ranked by median fitness. Note fitness is displayed on a logarithmic scale to emphasize relative differences in fitness between variants. Boxplots show median and exclusive interquartile range.

### Longitudinal trajectories allow for a more accurate stratification of CHIP variants

Since longitudinal data allow direct quantification of the growth in VAF over time, we can inspect the gradients (fluctuations) in VAF for variants that were classified as CHIP based on thresholding. We find that a VAF>0.02 threshold not only misses fast growing and potentially harmful variants (Fig.2A), but can include variants whose frequencies are shrinking (Fig.2A and 2B) and thus either do not confer a fitness advantage or are being outcompeted by other clones. Overall, only 56% of CHIP mutations detected by thresholding at 0.02 VAF were growing during the observed time span (Fig.2B). Longitudinal data thus reveal limitations in defining CHIP mutations based on a widely used VAF threshold. We therefore sought to filter variants based on longitudinal information, by selecting variants whose growth cannot be explained by the drift in VAF from neutral fluctuations (Fig.3A). Using the longitudinal behaviour of synonymous mutations in the LBCs, as a proxy for neutral trajectories, we find that the predicted linear relationships for mean and variance of VAF-gradients (see Methods) accurately reflect the data (Fig.3B). This novel approach, which we named drift-induced fluctuations (DIF) filter, allows us to estimate the distribution of fluctuations due to the neutral drift of clone populations as a function of VAF and detects variants growing more than two standard deviations from the mean of the predicted neutral drift (Fig.3A, Table7). DIF-filtering resulted in 226 variant trajectories (Fig.3C), 97% of which grew over the observed time span. We note that the VAF of fit mutations may still shrink over time due to the presence of an even fitter clone in the same individual. This is in contrast to thresholding at 0.02 VAF, with only 56% of variants identified to be growing and thus likely to confer a fitness advantage. Of 226 variants we detected, only 81 would have been detected using the previous VAF-threshold filter approach. We recomputed fitness estimates for this new set of filtered trajectories (Fig.3F). Growing variants that were missed by the traditional filtering method include highly fit variants such as *U2AF1* c.470A>G and *NPM1* c.847-4C>T (Fig.3D and 3F). VAF-thresholding identified *NOTCH1* as one of the top three most frequently mutated genes in our cohort. However, only 1 of the 8 variants identified was growing over the observed time-course. In contrast, all 6 *NOTCH1* variants identified by using our DIF filter were growing, thus our filtering method allows us to more effectively separate benign from potentially harmful variants. Overall, the variants only detected by DIF-filtering, but not VAF-thresholding, were of high fitness (Fig.3E).

**Figure 3.**
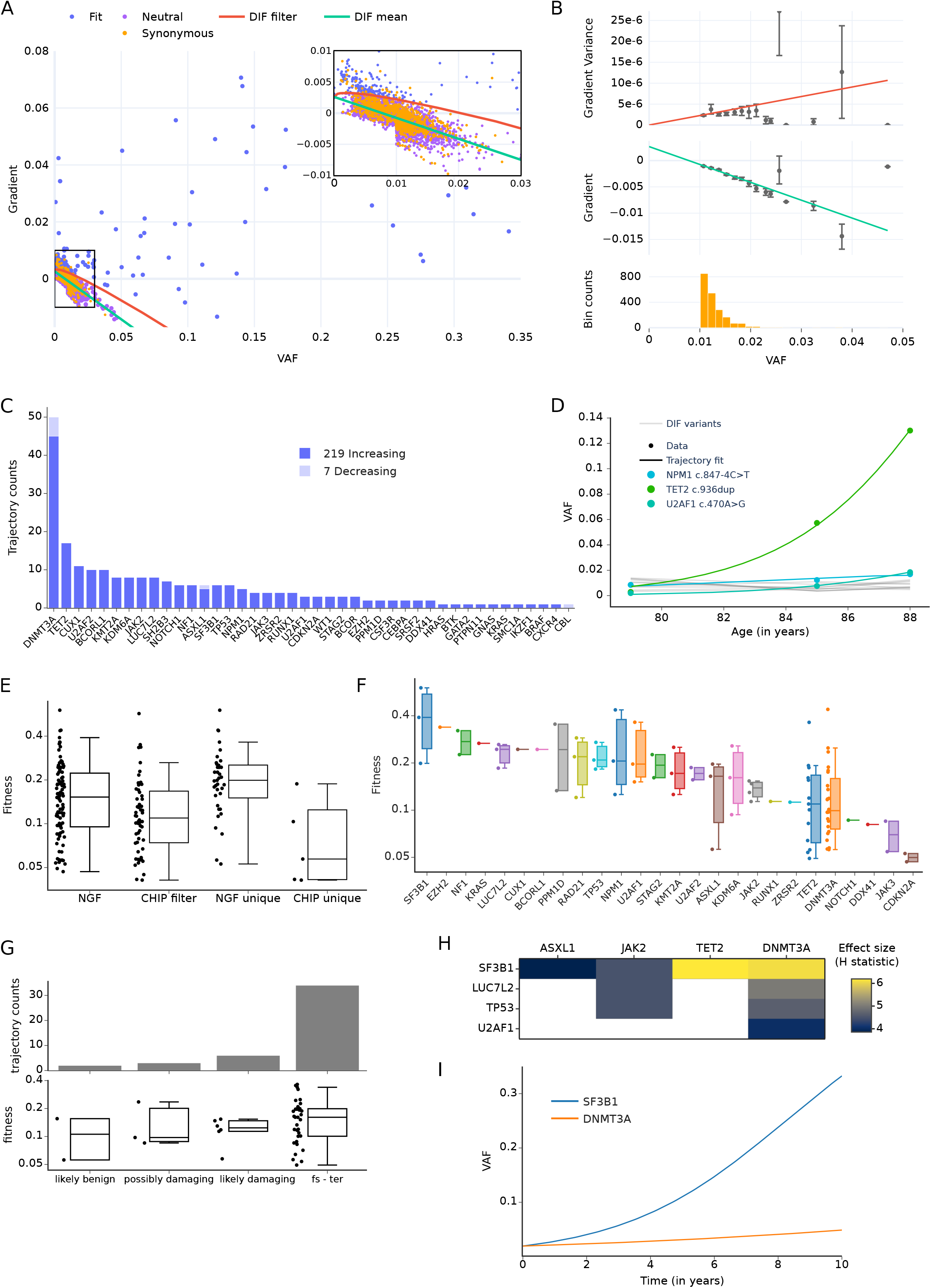
Filtering trajectories based on whether they exceed neutral fluctuations. A. Time-normalised gradient in VAF vs VAF for variants detected in the LBC with at least 2 time-points and at least one VAF >0.01 per trajectory, with synonymous (orange dots), non-synonymous fit (blue dots) and likely neutral (purple dots) mutations, classified based on whether the gradient exceeds the drift-induced fluctuations (DIF) of synonymous mutations (red line, two standard deviations above mean, green line). Inset: Close-up of low VAF region. Note that data below 0.01 VAF were only included if the trajectory later rises above 0.01, hence the data are less dense below VAF=0.01. B. Statistics of fluctuations of synonymous mutations. Both the mean (middle) and variance (top) of time-normalised VAF-gradients were fitted with a linear model. Only data at VAF ≥ 0.01 where used as data below this are incompletely sampled (see A). The fit was weighted by the number of data in each bin (bottom). C. Number of trajectories passing the DIF filter, broken down into whether VAF is increasing or decreasing from first to last time-point. D. Variant trajectories in one individual classified according to the DIF filter. Mutations with positive fitness effects are coloured according to variant classification (see Fig.1B) while likely neutral trajectories are shown in grey. For fit variants, points correspond to the observed VAF measurements and lines to the model fit. While the TET2 variant was also detected using the 0.02 VAF-threshold, the NPM1 and U2AF1 were only detected using the DIF-filter, as they did not reach 0.02 VAF. In particular the U2AF1 variant is fast growing (s=0.364). E. Fitness effects of variants broken down by filtering method. F. Fitness effects of mutations ranked by median fitness. Note fitness is displayed on a logarithmic scale to emphasize relative differences in fitness between variants (E&F). Boxplots show median and exclusive interquartile range G. Fitness effects of variants broken down by predicted mutation effect (see Methods). H. Analysis of variance of the distribution of fitness across genes. Heatmap of all statistically significant (p<0.05) Kruskal-Wallis H statistics, labeled by effect size, computed for all combinations of pairs of genes. I. Predicted growth of SF3B1 and DNMT3A variants over 10 years, all starting at 0.02 VAF, using the median fitness effects per gene (F).

We further stratified variants using a combination of computational predictors (Fig.3G, see Methods), categorising the most prevalent CHIP variants into damaging (6 variants), possibly damaging (3 variants) and likely benign (2 variants), as well as frameshifts and terminations (34 variants, which are also most likely damaging to protein structure and thus protein function). The novel DIF-filter therefore produces a low false discovery rate of pathogenic variants, with 88% of the detected fit variants being predicted to be likely damaging, frameshift or termination. As our approach detects very few benign variants by design, we were unable to report statistically significant differences between the fitness distributions classified by damaging. Further, predicted benign mutations selected as fit could be the result of clonal hitchhiking (mutations co-occurring in cells carrying a damaging mutation).

Taken together, measuring changes in synonymous variants combined with mathematical models of neutral drift results in an alternative method, DIF-filtering, that improves on the threshold-based definition of CHIP mutations (Fig.3A), by replacing an arbitrary cut-off on VAF by a choice of false discovery rate and as a result selecting fewer trajectories with shrinking VAF (Fig.2B and 3C). Importantly, only 2 time points are necessary to apply DIF-filtering, making this a widely applicable method for existing cohorts and future studies.

### Predicting trajectories using our fitness estimates from single-time point data

We further analysed differences in the distributions of fitness between genes using a non-parametric test. Despite having small sample sizes for many genes we still detected statistically significant differences among the distributions of fitness effects (Fig.3H). In particular, we found that mutations in gene *SF3B1* conferred a higher fitness advantage over mutations in common genes such as *DNMT3A, TET2* or *ASXL1*. Differences in the distribution of fitness allow us to predict the future growth of mutations from initial time-points. For example, if a patient presents with a variant in gene *SF3B1* or *DNMT3A* at 0.02 VAF, their growth is predicted to differ by 158% percent in 3 years (respectively achieving 0.057 and 0.026 VAF, assuming median fitness for each gene), warranting a clinical follow-up over that time-frame. We have also tested differences in fitness by genes when summarised into functional categories and found trajectories of genes involved in DNA methylation to have lower fitness than genes involved in splicing, the cohesin complex and histone methylation (SuppFig.2G and 2H).

## Discussion

The clinical potential for stratifying progression of CHIP depends on whether genes confer distinct fitness advantages. Indeed, most studies so far have not shown a clear distinction of fitness effects on a gene basis, and have shown considerable overlap in fitness coefficients between variants of different genes. We show that fitness can substantially differ by gene and gene category. Combining longitudinal data with a new method to identify CHIP variants allows for more accurate fitness estimates of CHIP than cross sectional cohort data, and motivates further studies with increased sample sizes.

The strength of our approach, combining longitudinal data with our new DIF-filtering method, is exemplified by *NOTCH1* and *LUC7L2*. *NOTCH1* variants identified by a 0.02 VAF threshold were prominent in the LBCs (Fig.1A). However, all but one of these were shrinking in frequency. In contrast, our DIF-filtering method identified 6 variants conferring a fitness advantage, including the only increasing variant detected at a 0.02 VAF level, c.4598A>C (Fig.3C). Although *NOTCH1* mutations commonly occur in leukaemia, this specific mutation has not been reported in COSMIC, thus more cohort studies are needed to clarify a role for *NOTCH1* in CHIP (Ferrando, 2009; Rossi et al., 2012). *LUC7L2* is a pre-mRNA splicing factor that has been previously reported to be involved in CHIP, MDS and AML (Hershberger et al., 2016; Sperling et al., 2017). *LUC7L2* was excluded from our list of CHIP mutations when using a 0.02 VAF cut-off. In contrast, we find several *LUC7L2* mutations using DIF-filtering, thus supporting previous findings. Other well-described CHIP genes that were only detected using DIF-filtering include *NPM1, TP53* and *U2AF2* (see Fig.2E and 3F,(Jaiswal et al., 2014)). DIF-filtering also results in higher fitness estimates for some genes, such as *PPM1D, U2AF1* and *RAD21*, because we either detected more trajectories of a gene with the DIF filter or were able to detect other variants at small VAF in the same individuals (Fig.2E and 3F). Importantly, a minimum of two time points per participant is needed to apply DIF-filtering, making this method widely applicable to future prospective studies and existing cohorts.

Synonymous mutations in our study reached a VAF of up to 0.02 (Fig.3B). Some previous studies have claimed that neutral events will typically not expand to > 0.001 VAF over a human lifespan (Lee-Six et al., 2018), and that all synonymous variants detected are likely hitchhikers (Watson et al., 2020). Our longitudinal data clearly show many synonymous and non-synonymous mutations larger than 0.001 VAF that do not grow in VAF (Fig.3A) or whose growth and shrinkage over time is consistent with neutral drift (Fig.3B and SI methods). If the synonymous mutations were able to grow to such high frequencies by hitchhiking on clones with fit mutations, these no longer have a fitness advantage over the observation period. An alternative explanation considers the number N of self-renewing cells (HSPCs). For an individual aged *t*, neutrally drifting mutations are only expected to achieve 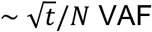. Recent studies, therefore, suggest that the high frequency at which CHIP is found in the elderly is likely the result of a small pool of stem cells (Zink et al., 2017). Estimates for the number of HSPCs in healthy adult humans vary widely, with recent phylogenetic estimates suggesting 50k-200k (Lee-Six et al., 2018), whereas other estimates suggest 11-22k (Abkowitz et al., 2002). On the lower end, a study based on allometric scaling estimates that the “pool of actively replicating cells that contribute to hematopoiesis” is around 500 cells (Dingli and Pacheco, 2006). In our study, the distribution of gradients of synonymous variants provides the estimate *θ* ~ *λ/N*, where *θ* corresponds to the slope of the linear relation between variance and vaf (Fig.3B) and *λ* to the rate of self-renewing divisions. Using the well-established rate *λ* = 1.3/year (once every 40 weeks, (Catlin et al., 2011)) we obtain an estimate on the number of HSPCs of *N* ~ 6500 (see SI methods for a more detailed analysis). The presence of neutral events reaching large VAFs in our data could be explained by a depletion of the stem cell pool in participants of the LBC (80 years mean across all collected blood samples). Both a study of somatic mutations in a 115-yr-old woman and a recent longitudinal study following the evolution of clones in an individual from age 103 to 110 (3 time points) found that likely benign clones achieved VAFs of up to 0.3 (van den Akker et al., 2020; Holstege et al., 2014)-suggesting a very small pool of stem cells. Another possible explanation is that the observed variants arose in cells with faster rates of self-renewing symmetric divisions *λ*, either due to natural selection of faster dividing HSPCs or because the observed variants in the LBC arose in different HSPC compartments, such as multipotent progenitor cells (Barile et al., 2020; Morcos et al., 2020; Takahashi et al., 2021). Since we can only directly measure *θ*, it is possible that both a depletion of the pool of HSPCs at old age and heterogeneity in the self-renewal rate *λ* have enabled neutral events to reach high VAFs. Despite the disparity of estimates on the number of HSPCs in healthy individuals between studies, the fitness estimates we report (Fig.2 and Fig.3) are valid for the wide ranges of reported HSPC estimates (SuppFig.2E and 2F and SI methods).

In this work, we have assumed each mutation to uniquely identify a clone. It is possible for multiple mutations to have occurred in the same clone, though this has been estimated to be as rare as 0.16 double mutants per person, for mutations conveying a fitness advantage *s* = 0.1, achieving VAF > 0.005 at age 80 (Watson et al., 2020). There may be a few variants that are double mutants, and these would increase the distribution of fitness reported for individual genes (Fig.2C and 3F), but not strongly affect the median fitness per gene. Double mutant clones in which one mutation is a hitchhiker contributing little to the fitness effect could be identified by seeking trajectories with very similar fitness values. Further work could estimate the fitness effects of double mutants by identifying possible subclone structures and then fitting models separately for each case. Ultimately, we will need single-cell targeted sequencing or colony growth essays combined with sequencing to verify the identity of clones with multiple mutations.

## Acknowledgments and author contributions

We thank Prof. Chris P. Ponting for critical reading of the manuscript.

The LBC1921 was supported by the UK’s Biotechnology and Biological Sciences Research Council (BBSRC), a Royal Society–Wolfson Research Merit Award to I.J.D., and the Chief Scientist Office (CSO) of the Scottish Government’s Health Directorates. The LBC1936 is supported by Age UK (Disconnected Mind project, which supports S.E.H), the Medical Research Council (MR/M01311/1, MR/K026992/1, which supported I.J.D.), and the University of Edinburgh. KK is funded by a John Goldman Fellowship sponsored by Leukaemia U.K. (2019/JGF/003). M.T.T. and N.R. are supported by MRC funded Ph.D. studentships (MR/N013166/1). TC and LS are supported by Chancellor’s Fellowships held at the University of Edinburgh. JAM is a Lister Institute Research Fellow. ELC is a cross-disciplinary post-doctoral fellow supported by funding from the University of Edinburgh and Medical Research Council (MC_UU_00009/2). We are grateful for funding from the Howat Foundation. LS, KK and TC conceived and supervised the study. NAR, ELC, LS, KK and TC wrote the manuscript. LM, AF, LMG generated data. NAR and ELC developed the methodology for data analysis. NAR, ELC, MTT, ACP, JAM, BJL conducted data analysis. SEH, SC, IJD curated the LBCs and gave access to samples and data. MC advised on aspects of the study.

## Data Availability

Data are deposited in the European Genome-Phenome Archive and provided via the LBC DAC (https://www.ed.ac.uk/lothian-birth-cohorts/data-access-collaboration)

## Supplemental Tables

Supplemental tables are available at the following link: https://drive.google.com/file/d/10UMVtRk8rKosBJE3fjnOMpbYyrvgjD_K/view?usp=sharing

## Supplemental Figure Legends

**Figure S1.**
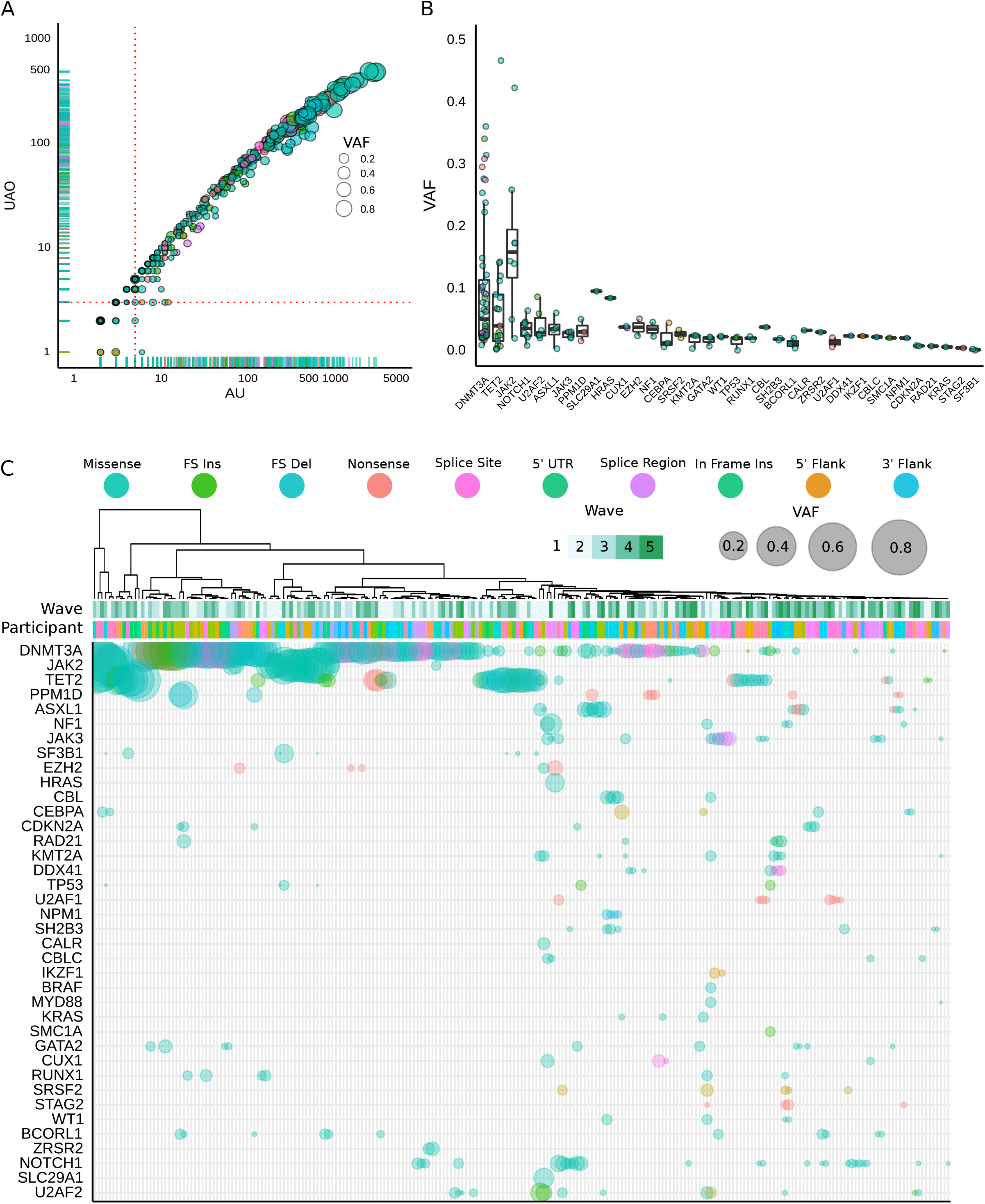
Clonal Haematopoiesis in the Lothian Birth Cohorts. A. Sequence quality metrics for our mutation calls across all participants and time-points filtered for 0.02 VAF. Plotted here are the AO (the number of sequenced reads supporting the alternative allele (mutation)) against the UAO (the number of sequenced reads with unique start sites that support the alternative allele - an additional measure of molecular complexity in our sequencing libraries). The red dotted lines denote the filter thresholds in both measurements (AO ≥ 5, UAO ≥ 3) and points are scaled by the VAF of the somatic mutation. Having mined for participant matched data points for any variant that surpasses our 0.02 VAF threshold, only 12 data points failed to meet our filter criteria (of 466). We did not exclude these as they were supported with matching events across any participants’ time series. B. Box and jitter plot of the variant allele frequency of all observed events in the 1st Wave at a 0.02 VAF filter coloured by variant classification and ordered by largest mean VAF. C. Schematic of all affected genes in the cohort with the largest clone size of an event in any given gene shown. All affected participants have been clustered across all time-points, with the point size scaled by VAF and coloured by the functional consequence of the variant (as per legend).

**Figure S2.**
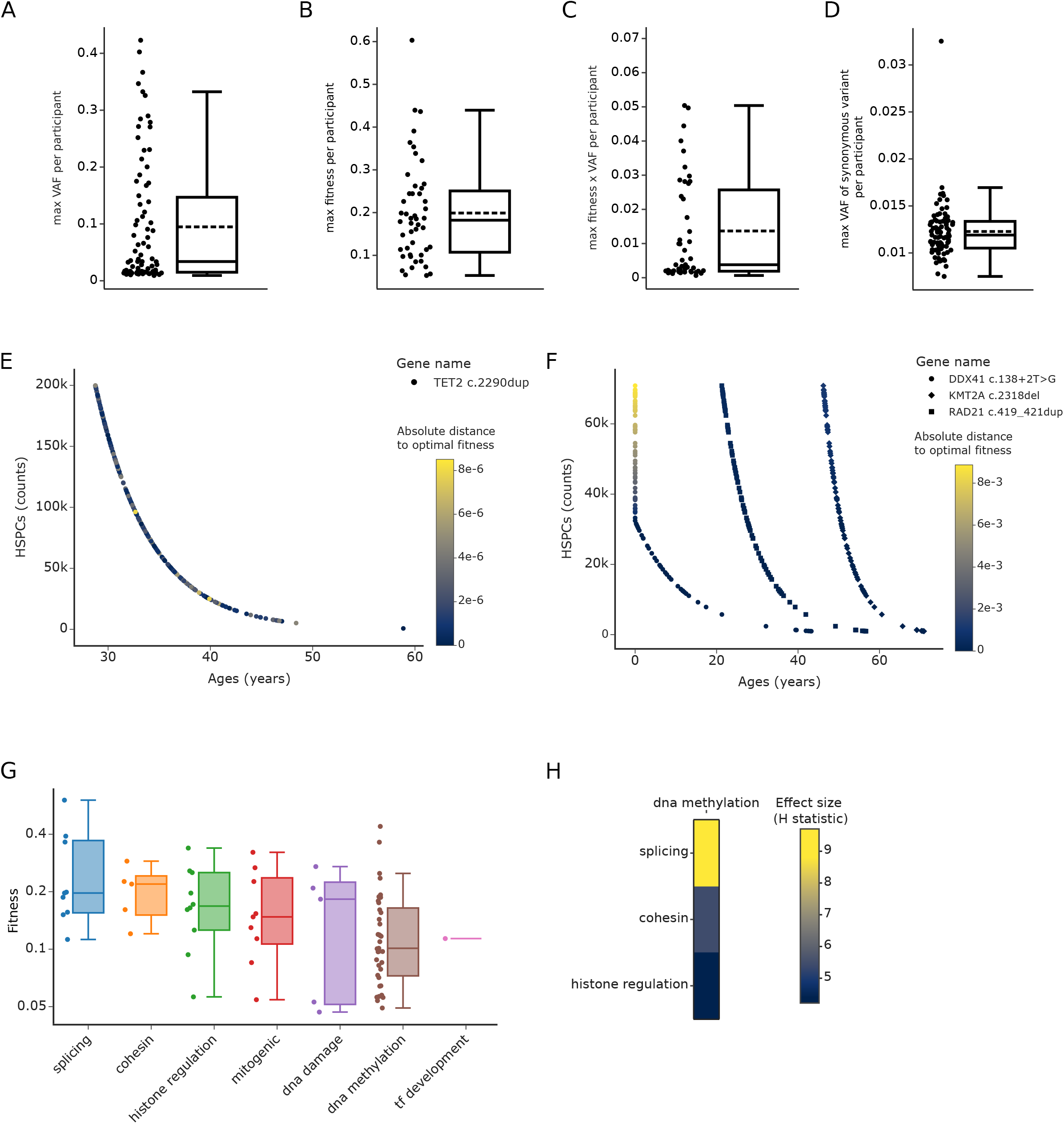
Cohort properties for fitness estimates. A - D. Box plots show median (bold line) and mean (dashed line) and exclusive interquartile range. A. Distribution of maximum VAF, *v*, per participant of the LBC. *v* is computed as the mean over all observations of a trajectory. One trajectory with *v* > 0.5 was excluded for this distribution. B. Distribution of maximum inferred fitness, *s*, per participant of the LBC. This distribution only shows participants with at least one trajectory selected as fit using the DIF filter. C. Distribution of maximum inferred *s* * *v* per participant in the LBC. *s* corresponds to the inferred fitness of a variant and *v* to its average observed VAF. This distribution only shows participants with at least one trajectory selected as fit using the DIF filter. D. Distribution of maximum VAF, *v*, of synonymous mutations per participant of the LBC. *v* is computed as the mean over all observations of a trajectory. E. Hypersurface of parameters producing the same fit to the evolution of a fit clone in a participant of the LBC. Each marker corresponds to a combination of parameters (fitness, number of wild type HSPCs in individual and time of mutation acquisition) fitted to the trajectory of a participant in the LBC. The colour scale shows the absolute difference between the inferred fitnesses associated with each marker and the fit producing the least squared error. All markers shown are within a relative tolerance of 10^−3^ of the solution presenting the least squared error. F. Hypersurface of parameters producing similar fits to the evolution of clones in a participant of the LBC with 3 fit variants. Each marker corresponds to a combination of parameters (fitness, number of wild type HSPCs in individual and time of mutation acquisition). Curves were fitted simultaneously, sharing a common number of HSPCs. The colour scale shows, for each variant, the absolute difference between the inferred fitnesses associated each marker and to the fit producing the least squared error. All markers shown are within a relative tolerance of 2×10^−3^ of the solution presenting the least squared error. Notice that the fitness of variant *DDX41* marginally increases when the time of mutation acquisition meets the boundary at age 0 in the exponentially decreasing hypersurface of parameter solutions. G. Fitness effects of mutations grouped by gene category and ranked by median fitness. Note fitness is displayed on a logarithmic scale to emphasize relative differences in fitness between variants. H. Analysis of variance of the distribution of fitness across genes. Heatmap of all statistically significant (p<0.05) Kruskal-Wallis H statistics, labelled by effect size, computed for all combinations of pairs of genes.

## Supplementary Methods

### 1 Birth-death model of clonal dynamics

Consider a population *X*(*t*) of haematopoietic stem and progenitor cells (HSPCs) in an individual at time *t*. A classical model of stem cell population assumes that HSPCs divide and differentiate according to the following birth-death process, (Till et al., 1964; Watson et al., 2020):

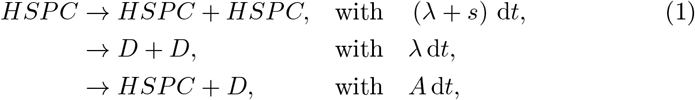

where D denotes a differentiated cell. Since differentiated cells cannot self-renew, differentiation will result in cell death and is treated as such. The deterministic behaviour of this system is given by

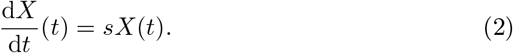

Consequently, the time evolution of the population of HSPCs is

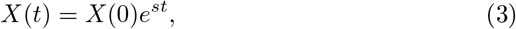

where *X*(0) corresponds to the initial population at time *t* = 0. It is clear that parameter *s* regulates the excess growth towards self-renewal and dictates the evolution of *X* (*t*).

The stochastic behaviour of this model, on the other hand, is more involved. Assuming that *X* (0) = 1, *X* (*t*) has the following probability distribution (Bailey, 1990):

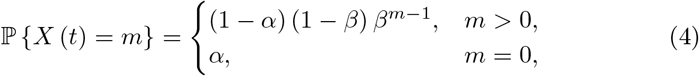

where

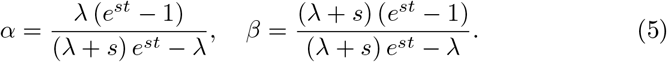

More generally, the distribution accounting for the case when the initial population is *a >* 0,

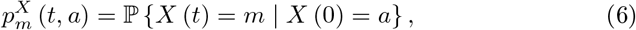

can be analytically derived as

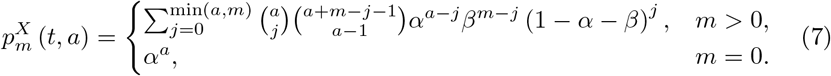

Of particular relevance is the mean and variance of *X* (*t*), if *X* (0) = *a*:

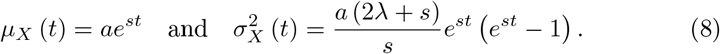

### 2 A stochastic model of neutral clones

Assume that all HSPCs in an individual follow a critical birth-death (CBD) process (*s* = 0). When a mutation occurs in a HSPC it gives rise to a genetic clone *c_i_* with

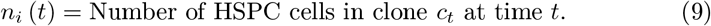

For now, we assume that mutations have no impact in the bias towards self-renewal, *s*, and that *n_i_* in turn follows a CBD process. We refer to 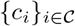 as the collection of all *neutral clones* (*s* = 0) present in the individual. If we denote by *t_i_* the time of acquisition of the mutation, so that *n_i_* (*t_i_*) = 1, then the probability distribution of *n_i_* (*t*) is given by the limiting case *s* → 0 of (4), (Bailey, 1990). That is, for any *t > t_i_*,

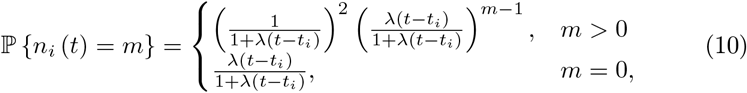

Further, given *t*_0_ *> t_i_*, for any *t > t*_0_ consider

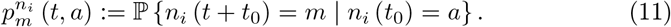

The mean and variance of this probability distribution can be derived, again, as the limit *s* → 0 of (8),

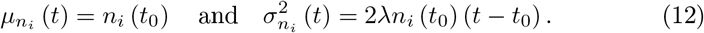

#### 2.1 Distribution of VAF gradients in neutral clones

Next, we want to understand how the stochastic evolution of HSPC counts over time translates to the evolution of the variant allele frequency (VAF or blood share) of genetic clones. Further, rather than the distribution of VAFs at a particular time, we want to derive the distribution of expected gradients of VAF between two time points.

##### 2.1.1 Linear scaling of random variables

First, let us remind the scaling properties of the mean and variance of a random variable under affine transformations. Let *X* be a random variable with mean *μ_X_* and variance 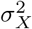 and define *Y* as

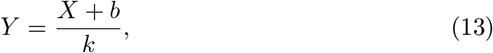

for scalars *b* and *k*. Then mean and variance scale as follows,

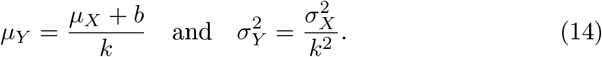

##### 2.1.2 Distribution of VAF evolution

Since we assume that all HSPCs in the individual follow a CBD process, the total population of HSPCs remains stable, on average, at *N*. Further, since asymmetric divisions are far more common than symmetric divisions, we can assume that the number of differentiated blood cells produced by a clone of HSPCs is directly proportional to the clone’s size. We therefore model the VAF of a clone *c_i_* at time *t > t_i_* as

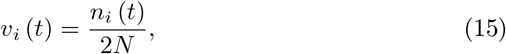

where the factor 2 in the denominator is the result of diploidy in HSPCs.

Further, for any time *t > t*_0_ *> t_i_* we can consider the probability distribution 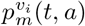 of the evolution of VAF from an initial state *v_i_*(*t*_0_) = *a* as in (11). It follows from the distribution of clone sizes (12) and scaling properties (14) that

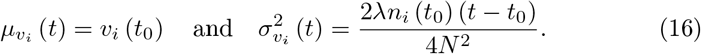

When observing VAF distributions, it is worth noting the natural emergence of parameter

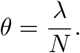

A CBD model of the evolution of clones in HSPCs, therefore, yields that the variance of the distribution of VAFs at time *t* from an initial state *v_i_*(*t*_0_) is

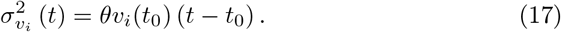

##### 2.1.3 Distribution of VAF gradients between two time points

Again, for any *t > t*_0_ *> t_i_* consider the VAF evolution *v_i_*(*t*) of a neutral clone *c_i_* and the resulting probability distribution of the time-adjusted gradient

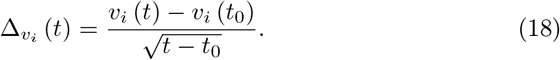

From scaling properties (14) and the distribution of VAFs of neutral clones (16) and (17), it follows that

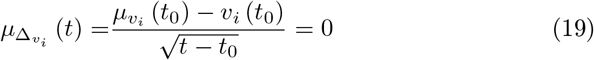

and

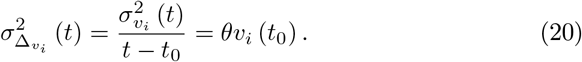

Notice that by considering a time-adjusted gradient (taking the square root of the time difference) the variance of 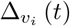 becomes independent of time.

### 3 A stochastic model of neutral clones in the presence of a fit clone

In the following we want to study the evolution of a neutral clone’s gradient in the presence of a non-stochastic growing mutation (the deterministic limit for *s* ≫ 0). Consider an individual with a collection of neutral clones, {*c_i_*}_*i*∈*I*_, that harbours a mutation at time *t_f_* giving rise to a clone *c_f_* with a fitness advantage *s >* 0.

In the continuum limit, the deterministic evolution (3) of the clone’s size *n_f_*(*t*) is

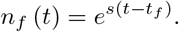

The total pool of HSPCs in this individual is, therefore, comprised of a collection of wild type HSPCs that remains constant in time at *N_w_* and a mutant growing population of size *n_f_*(*t*):

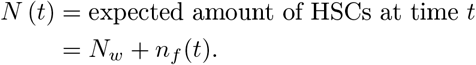

The evolution of the time-adjusted VAF gradient of neutral clones then becomes

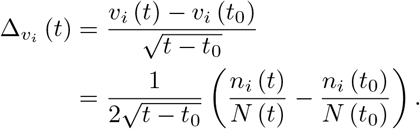

#### 3.1 Neutral drift in the presence of a fit clone

Here we will show that 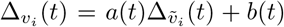, where 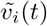 corresponds to the evolution of a neutral clone’s VAF in the absence of fit clones,

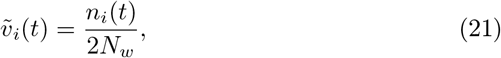

and *a*(*t*) and *b*(*t*) are functions of time, independent of the random process *n_i_*(*t*).

First, notice that

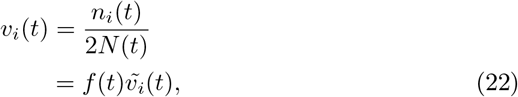

with *f* (*t*) = *N_w_/N* (*t*). The mean value theorem asserts that ∃*ξ* ∈ [*t*_0_*, t*] so that

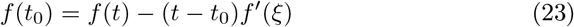

and

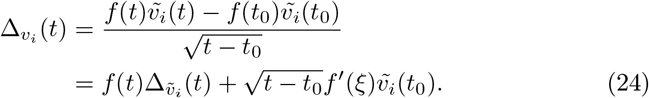

#### 3.2 Estimating the mean gradient of neutral variants

Since the expectation operator *μ*_(·)_(*t*) is a linear operator and 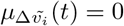, (24) yields

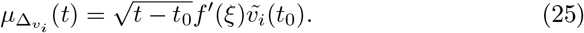

From the definition of *f*(*t*) it follows that, *f*′(*ξ*) = 2*sv_f_*(*ξ*)*f*(*ξ*). Combining this expression with (22) we have

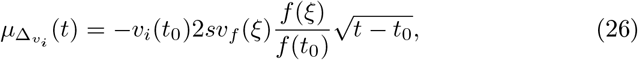

and assuming *f*(*ξ*) ≈ *f*(*t*_0_),

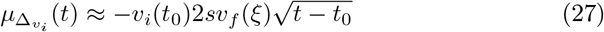

for *ξ* ∈ [*t*_0_*, t*]. We use the Lothian Birth cohort (LBC) data to extract statistics about the distributions of VAF and fitness of clones to check that this linear relation holds true (Fig.3B and SuppFig.2A-D). We find that the median over all participants with a measurable fit mutation (fit mutations can occur outside the set of sequenced genes) of the maximum realisation of *v_f_* (*t*) _*_ *s* across all present fit clones is

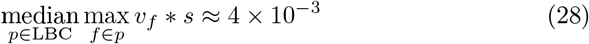

where *v_f_* is the averaged VAF over all data points (SuppFig.2C). Further, the average over all combinations of *t, t*_0_ in our data is 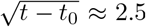. With these we estimate

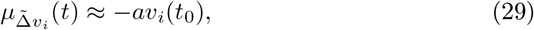

with *a* ≈ 0.02.

We estimate the mean of VAF fluctuations of neutral clones by averaging the fluctuations of synonymous trajectories (clonal trajectories of genetic variants not changing the coded amino acid) with the same VAF, *v*, across all participants, 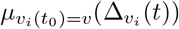. Fitting a weighted linear regression model on our data yields

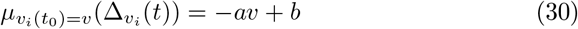

with *a* ≈ 0.3 and an intercept *b* ≈ 10^−3^ to account for the lack of observations of extinction events (Fig.3B). There are a number of possible reasons for the numerical discrepancy of the theoretically estimated and empirically observed slope, such as the approximation *f* (*ξ*) ≈ *f* (*t*_0_) made above, but also the fact that many individuals will have multiple fit clones affecting the gradients of VAF of neutral variants.

#### 3.3 Estimating the variance of gradients

From (24) and scaling of random variables (14) we find

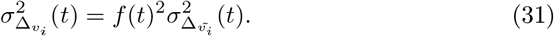

First notice that, in the presence of a small fit clone *n_f_*(*t*) *< N_w_ ⇔ v_f_* (*t*) *<* 0.25, the scaling factor*f* (*ξ*) is bounded by 1/2 *< f* (*ξ*)^−1^ *<* 1. This initial approximation yields a bound on the variance of fluctuations in the presence of small clones:

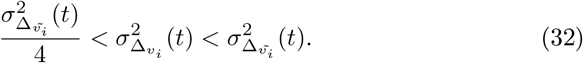

A more careful analysis shows a linear relation 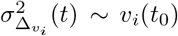. Combining (31) and (17) we obtain

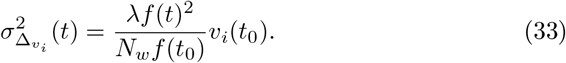

Then, approximating *f* (*t*) ≈ *f* (*t*_0_) yields

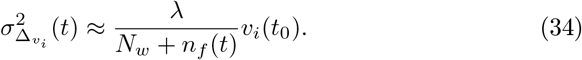

Finally, from the definition of VAF it follows that 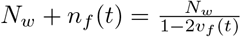 and

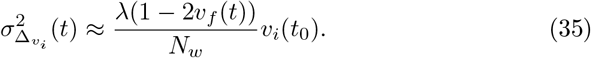

To estimate 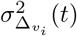 from the LBC data we approximate 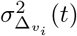 by the variance of observed gradients over all synonymous trajectories with *v_i_*(*t*_0_) = *v*, 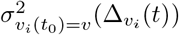. To compute the variance over trajectories with *v_i_*(*t*) = *v* we bin trajectories by VAF size. Then a linear regression fit using weighted least squares (weighted by bin size) yields (Fig.3B):

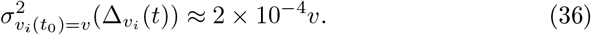

### 4 Estimation of the number of HSPCs in participants of the LBC

Combining (35) and (36) we can provide an approximation of the number of HSPCs in an individual

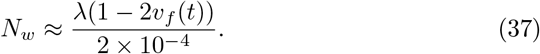

Denoting by *v_i_* the mean VAF of a clone *v_i_*(*t*) over the observed time span in the LBC, we approximate

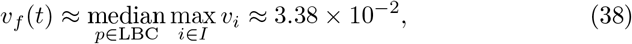

as the median over all participants in the LBC of the mean VAF of their largest clone (SuppFig.2A). Further, we use the well-established estimate that HSPCs divide symmetrically for self-renewal every 40 weeks (Catlin et al., 2011), *λ* = 1.3, and estimate

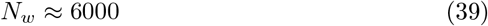

#### 4.1 Estimations not accounting for the presence of fit clones

Since fit variants remain relatively small, notice that estimate (39) does not differ much from the estimate obtained directly from the evolution of neutral clones in the absence of a fit clone (20),

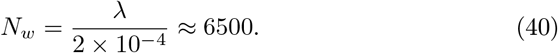

An even simpler approach to estimating the number of HSCs is provided by the distribution of neutral clones (16). At age *t* the maximum VAF reached by neutral clones, *c_i_*, should be.

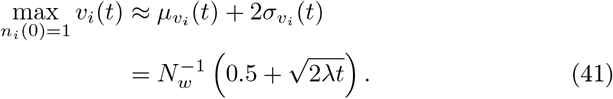

In the LBC data, we estimate 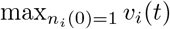 using the mean size and age of synonymous trajectories *v_s_* (SuppFig.2D):

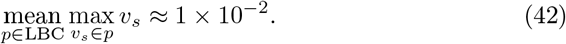

Using the average age of participants during the observation of the trajectories used to compute the above estimate, *t* ≈ 80 years, we obtain the rough estimate

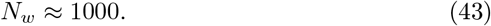

### 5 Drift-induced fluctuations filter

In the LBC, we estimate linear relations (30) and (36) and select clonal trajectories, *v*(*t*), growing more than the expected growth over the observed time span,

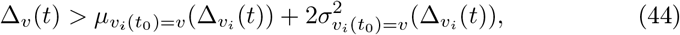

as mutations conferring a fitness advantage, *s >* 0.

Notice that, although this sieving method depends on the length of the observation period *t − t*_0_, estimate (30) takes into account the average observation period of clonal trajectories. Further, the square root scaling of time in (27) reduces in the impact of the difference of observation periods; a span of 3 to 5 data points per trajectory yield 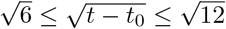.

### 6 Estimating clonal fitness

Given an individual with a collection of fit clones {*c_i_*}_*i*∈*F*_ with fitness parameters {*s_i_*}_*i*∈*F*_, their predicted deterministic time evolution is

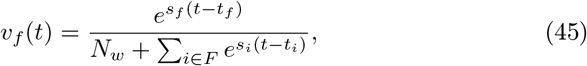

for *f* ∈ *F*. We fit this evolution to data using least squares to infer parameters (*s_f_, t_f_*) for each clone {*c_f_}_f_*_∈*F*_ and a parameter *N_w_* shared across all clones present in the individual.

We note the existence of a hypersurface of parameters yielding the same evolution. In the case of a single fit clone, *v_f_*(*t*), equation (45) can be re-parametrised as

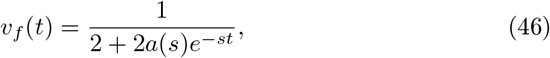

with 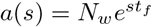. Therefore, any combination of parameters *N_w_* and *t_f_* satisfying 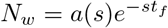 will produce the same VAF evolution *v_f_* (*t*) (SuppFig.2E and 2F). Let us highlight that, although different choices of *N_w_* will result in a different *t_f_*, the order of mutation acquisition remains unchanged with any choice of parameters in the specified hypersurface.

In this study we have refrained from fitting the 2-parameter model given both the impracticality of providing realistic bounds to *a*(*s*) (constraining *N_w_* within a pre-established ranges and *t*_0_ *< t_f_ < t* within achievable times as a function of fitness) and to show the existence of global solutions that span across different estimates of *N_w_* in the literature through no or minor adjustments of inferred fitness values (SuppFig.2E and 2F).

### 7 Multiple fit clones

If two clones with *s*_1,2_ *>* 0 are present in an individual, the VAF is given by

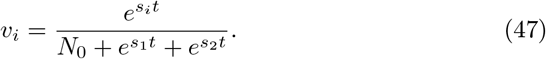

Assume without loss of generality that the second clone is fitter, *s*_2_ *> s*_1_, and consider under what conditions the VAF of the first clone will shrink, i.e.,

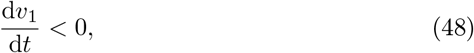

which is equivalent to

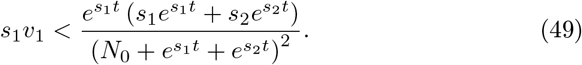

Solving for *t* gives

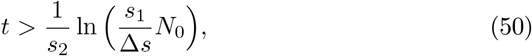

where Δ*s* = *s*_2_ − *s*_1_. From this we can see the fitter clone will always cause the first clone’s VAF to shrink for times greater than (50). As we expect, this happens earlier for fitter secondary clones (greater *s*_2_), and there is a further logarithmic dependence on the fitness of the first clone (later shrinking for greater *s*_1_), the size of the neutral HSPC pool (later shrinking for greater *N*_0_), and the difference in fitness between the two clones (earlier shrinking for larger Δ*s*).

